# Viral epidemic potential is not uniformly distributed across the bat phylogeny

**DOI:** 10.1101/2024.09.26.615197

**Authors:** Caroline A. Cummings, Amanda Vicente-Santos, Colin J. Carlson, Daniel J. Becker

## Abstract

Characterizing host–virus associations is critical due to the rising frequency of emerging infectious diseases originating from wildlife. Past analyses have evaluated zoonotic risk as binary, but virulence and transmissibility can vary dramatically. Recent work suggests bats harbor more viruses with high virulence in humans than other taxa. However, it remains unknown whether all bats harbor viruses of equal zoonotic potential. We used phylogenetic factorization to flexibly identify clades of mammals (at any taxonomic level) associated with low or high viral epidemic potential, and found virulence and transmissibility only cluster within bat subclades, often among cosmopolitan families. Mapping the geographic distributions of these bat clades with spatial data on anthropogenic footprint suggests high zoonotic risk in coastal South America, Southeast Asia, and equatorial Africa. Our results deepen understanding of the host– virus network and identify clades to prioritize for viral surveillance, risk mitigation, and future studies characterizing mechanisms of viral tolerance.

## Introduction

Due to the rising frequency of emerging infectious disease outbreaks in humans, there is an urgent need to characterize associations between viruses and their hosts ^1^. Over 70% of human emerging infectious diseases are caused by zoonotic pathogens, which originate in animals and adversely impact human health and economies ^1–3^. Statistical models can improve our capacity to preempt and mitigate zoonotic transmission by identifying the kinds of wildlife species most likely to harbor zoonotic pathogens ^4,5^. For example, past work has identified rodent and avian species likely to be undetected reservoirs of zoonotic pathogens ^6,7^. Species predicted to carry high-impact zoonotic pathogens can then be prioritized for surveillance and prevention measures^4,8,9^.

Zoonotic risk is often evaluated as a binary variable: whether a pathogen can or cannot infect humans ^5^. However, zoonotic pathogens differ in virulence (i.e., severity of disease) and transmissibility (i.e., capacity to spread in human populations after emergence)^8^. For example, within the genus *Betacoronavirus*, the case-fatality rates (CFRs) of viral species vary, with 5.9% in SARS-CoV and 34.4% in MERS-CoV ^5^. Further, while the former efficiently spreads from person to person, the latter is much less efficient at human-to-human transmission ^8,10^. To reduce zoonotic emergence and optimize time and resource allocation for public health measures, zoonotic pathogens with the highest virulence and transmissibility should be prioritized for surveillance and transmission prevention ^4,8^.

Recent work has demonstrated that bats (order: Chiroptera) harbor more viruses with high virulence in humans than other mammalian or avian orders ^8^. Bats are recognized as “special hosts” because they exhibit exceptional viral diversity (hosting more known viruses compared to most other taxa) and appear to tolerate many viruses without exhibiting clinical signs of infection ^11–14^. Bats hosting viruses with high virulence in humans can also be attributed to the evolutionary distance between us and these mammals ^8^. As the distance between a vertebrate host and humans increases, a virus is less likely to be pre-adapted to overcome human host defense mechanisms (e.g., immune function). However, humans are also less likely to be pre-adapted to cope with these novel infections and show resistance, increasing morbidity and risk of mortality ^5,15^.

As the only flying mammals, bats evolved diverse immune adaptations to cope with the metabolic demands of flight -that likely enabled them to tolerate otherwise virulent viruses ^11,16^. However, bats do not uniformly interact with viruses. As the second most speciose order of mammals (over 1,470 species)^17^, bats exhibit diverse ecological traits that may influence their viral communities, including their wide geographic distribution, use of torpor, heterogeneous diets, gregariousness, high mobility, long lifespan to body size ratio, and deep coevolutionary histories with viruses ^11,16^. Thus, the presumption that all bat species equally harbor a large number of virulent viruses may be inaccurate, and this has important implications for public health and bat conservation ^18^. For the latter point, bats have received negative media attention given their identity as reservoir hosts of several high-profile viruses (e.g., SARS-CoV, Nipah virus), leading to retaliation against bats ^16,19–21^. Clarifying the distribution of viral virulence across the order Chiroptera could provide the public with a more complete picture of zoonotic risk, thereby improving public perceptions of bats and informing outbreak prevention efforts^19,21^.

In this study, we test whether viruses with high virulence in humans are uniformly distributed across bats or if particular subclades of bats harbor more virulent viruses. We hypothesize virulence clusters within subclades of bats, based on the unique coevolutionary histories between different bats and viruses ^5^. First, to assess the phylogenetic distribution of viral epidemic potential, we quantify phylogenetic signal of CFRs and onward transmission across mammals and within bats specifically. Next, we use a flexible graph-partitioning algorithm to identify clades of mammals with unusually high or low viral epidemic potential without the need to *a priori* assume a specific phylogenetic scale ^22^. Finally, we mapped the geographic distributions of any identified bat clades with unusually high virulence in tandem with spatial data on anthropogenic footprint, visualizing hotspots of zoonotic risk to guide viral surveillance.

## Methods

### Viral epidemic potential data

We extracted mammal–virus associations from the Global Virome in One Network (VIRION), the most comprehensive-to-date database of vertebrate–virus associations ^23^. We retained only those mammals and viruses resolved by the National Center for Biotechnology Information (NCBI) taxonomy and matched mammals to the latest mammal phylogenetic tree using the *ape* package in R ^24,25^. There were 2,915 unique host–virus associations, of which 86% had a reported detection method (viral isolation, PCR, and/or serology; *n =* 2,507). Fifty percent (*n* = 1,459) represented detections through PCR (*n =* 1,103) and/or viral isolation (*n =* 797). Both methods can indicate active infection: viral isolation denotes infectious virus and thus successful replication in the host (i.e., implying competence and the capacity to infect new hosts), and PCR can detect the virus, but it may be noninfectious and thus non-transmissible ^26^. Only 5.5% of unique associations were detected solely using serology (*n =* 159), which is limited as it can signify either active infection or recent exposure ^27^. Further, serology can lack specificity because antibodies often cross-react with other pathogens ^27^. Finally, the remaining 14% of associations had no specified detection method (*n =* 408). To quantify typical viral virulence in humans attributed to each host, we averaged virulence across all viruses per host species using previously collated virulence data ^5^. We calculated the average and maximum CFRs, expressed as proportions of human cases resulting in mortality ^8^. We then calculated the typical transmissibility in humans attributed to each host as the fraction of viruses per host that have shown onward transmission in humans ^8^. Given the above mixture of viral detection methods, we caution that these data do not necessarily imply each host meaningfully contributes to zoonotic transmission of virulent viruses ^27,28^.

### Phylogenetic signal in viral epidemic potential

We first estimated phylogenetic signal for each response variable (mean CFR, maximum CFR, and onward transmission) using Pagel’s λ in the *caper* package ^29^. Pagel’s λ quantifies phylogenetic dependence in a trait by comparing observed variation to that expected under a Brownian motion model of evolution–traits gradually diverging over evolutionary time–with values of zero indicating complete phylogenetic randomness and values of one supporting high phylogenetic signal (i.e., Brownian motion) ^30,31^. We calculated λ across all viruses to test if viral epidemic potential is evolutionarily conserved across species in two analyses: among all mammals and within bats specifically. To assess phylogenetic patterns in viruses within well-studied viral groups, we repeated these analyses for viral families with sufficient sample size (≥ 30 host species)^32^.

### Taxonomic patterns in viral epidemic potential

We used a flexible graph-partitioning algorithm to identify clades of mammals and bats with unusually high or low viral epidemic potential. Phylogenetic factorization iteratively partitions a phylogeny to identify nodes with the maximum contrast in a given response variable, thereby identifying particular clades with greater or lesser propensities to harbor virulent or transmissible viruses without needing to *a priori* specify a given phylogenetic scale (e.g., mammalian order or family). We used the *phylofactor* package in R to partition each measure in a series of iterative GLMs for each edge in our phylogeny ^22^. We determined the number of statistically significant clades using Holm’s sequentially rejective test with a 5% family-wise error rate ^33^. We first performed phylogenetic factorization across all mammals to test if Chiroptera was organically identified as a clade with greater propensity to harbor virulent viruses. We next performed phylogenetic factorization within the order to identify particular bat clades with especially low or high viral epidemic potential in humans. As in our analyses of phylogenetic signal, we repeated these analyses for viral families with sufficient sample sizes.

### Geographic distribution of viral epidemic potential

In our analyses, we defined “risky” bat clades as those identified by phylogenetic factorization as harboring higher CFRs or propensity for onward transmission in humans. We aggregated the geographic ranges of bat species belonging to each risky clade using International Union for Conservation of Nature (IUCN) data ^34^. To visualize hotspots of potential viral epidemic burdens from bats, we co-mapped ^35^ anthropogenic footprint ^36^ and the geographic distributions of bats in risky clades to capture impacts on viral shedding and opportunities for cross-species transmission to humans.

## Results

### Phylogenetic signal in viral epidemic potential

Our data on viral epidemic potential included 983 mammal (220 chiropteran) species, infected by 115 unique virus species spanning 22 virus families. Mean CFR (x□ = 0.24 ± 0.01), maximum CFR (x□ = 0.39 ± 0.01), and fraction of viruses with onward transmission (x□ = 0.35 ± 0.01) all ranged from 0 to 1. Across all viruses and all mammals, mean and maximum CFR exhibited moderate phylogenetic signal (λ = 0.69 and 0.64, respectively), while the fraction of viruses with onward transmission exhibited high phylogenetic signal (λ = 0.85). All estimates of λ were statistically distinct from phylogenetic randomness (*p* < 0.001) and Brownian motion models of evolution (*p* < 0.001). Five viral families had sufficient sample sizes to enable stratifying our analyses: the *Coronaviridae* (*n =* 98 mammals), *Flaviviridae* (*n =* 381 mammals), *Rhabdoviridae* (*n =* 298 mammals), *Togaviridae* (*n =* 169 mammals), and *Paramyxoviridae* (*n =* 46 mammals).

Within each viral family, mean CFR showed high phylogenetic signal for coronaviruses; moderate phylogenetic signal for flaviviruses, rhabdoviruses, and togaviruses; and no phylogenetic signal for paramyxoviruses (Table S1, Figure 1). Maximum CFR exhibited high phylogenetic signal for coronaviruses, rhabdoviruses, and paramyxoviruses and moderate phylogenetic signal for flaviviruses and togaviruses (Table S1, Figure 1). Most λ estimates departed from both phylogenetic randomness (*p* < 0.001) and Brownian motion (*p* < 0.001), with two exceptions. Estimates of λ for mean and maximum CFR for coronaviruses did not depart from Brownian motion (*p* = 1), and mean CFR for paramyxoviruses did not depart from phylogenetic randomness (*p* = 1). The fraction of viruses with onward transmission showed high, moderate, and no phylogenetic signal for togaviruses, flaviviruses, and paramyxoviruses, respectively (Table S1, Figure 1). Togaviruses departed from phylogenetic randomness (*p* < 0.001) but not Brownian motion (*p* = 1). Paramyxoviruses departed from Brownian motion (*p* < 0.001) but not phylogenetic randomness (*p* = 1). Sample sizes for onward transmission were insufficient for the other two virus families.

**Figure 1.**
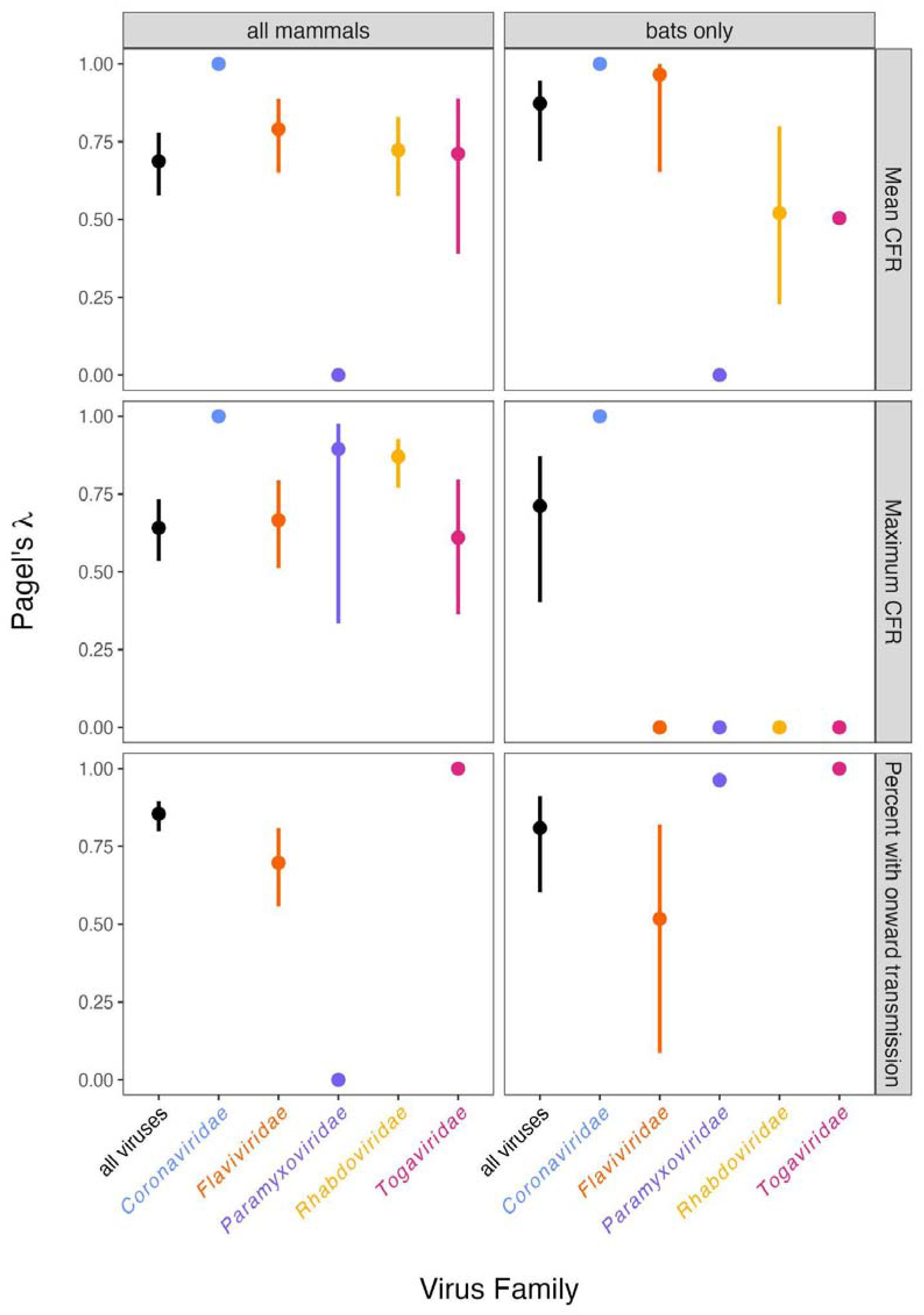
Phylogenetic signal (Pagel’s λ and 95% confidence interval) of mean CFR, maximum CFR, and the percent of viruses with onward transmission in mammal-wide and bat-specific analyses across all viruses and within the five virus families, with sufficient sample size. All λ estimates and tests of phylogenetic randomness or Brownian motion are provided in Table S1.

Within bats, mean CFR (x□ = 0.60 ± 0.03), maximum CFR (x□ = 0.81 ± 0.02), and the fraction of viruses with onward transmission (x□ = 0.32 ± 0.03) all ranged from 0 to 1. Across all viruses in bats, mean CFR and the fraction of viruses with onward transmission exhibited high phylogenetic signal (λ = 0.87 and 0.81, respectively), while maximum CFR showed slightly weaker phylogenetic signal (λ = 0.71). All λ estimates departed from both phylogenetic randomness (*p* < 0.001) and Brownian motion (*p* < 0.001).

Within each virus family in the bat-specific analysis, mean CFR showed high phylogenetic signal for coronaviruses and flaviviruses, moderate phylogenetic signal for rhabdoviruses and togaviruses, and no phylogenetic signal for paramyxoviruses (Table S1, Figure 1). Coronaviruses did not depart from Brownian motion (*p =* 1), and togaviruses and paramyxoviruses did not depart from phylogenetic randomness (*p =* 0.81 and 1, respectively). Maximum CFR showed high phylogenetic signal for coronaviruses, which departed from phylogenetic randomness (*p <* 0.001) but not Brownian motion (*p =* 1). The phylogenetic distribution of all other viruses did not depart from phylogenetic randomness (*p =* 1), and rhabdoviruses did not depart from Brownian motion (*p =* 1). The fraction of viruses with onward transmission showed high phylogenetic signal for togaviruses and paramyxoviruses and moderate phylogenetic signal for flaviviruses (Table S1, Figure 1); sample sizes for this measure were insufficient for the other two virus families. Togaviruses and flaviviruses departed from phylogenetic randomness (*p <* 0.001 and *p =* 0.03, respectively), and flaviviruses and paramyxoviruses departed from Brownian motion (*p <* 0.001). However, togaviruses did not depart from Brownian motion (*p =* 1), and paramyxoviruses did not depart from phylogenetic randomness (*p =* 0.26).

### Taxonomic patterns in viral epidemic potential

In the mammal-wide analysis, phylogenetic factorization identified 13 clades that differed in mean CFRs, 15 in maximum CFRs, and eight in the fraction of viruses with onward transmission (Figure 2, Table S2). Most of these clades (30/36 total, 9 of which were bat clades) were “risky” (i.e., harboring higher CFRs or propensity for onward transmission in humans compared to the rest of the host phylogeny).

**Figure 2.**
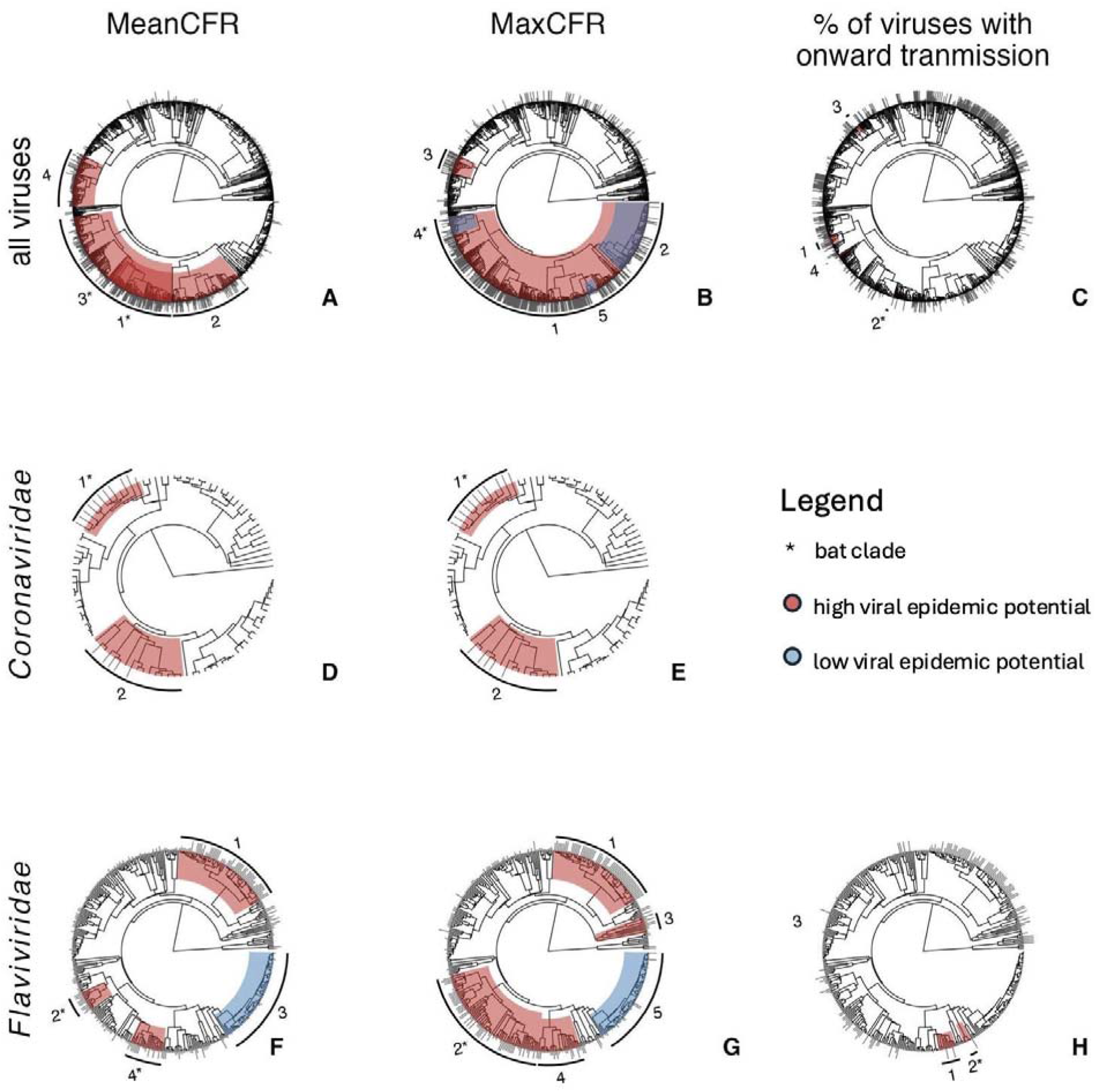
Phylogenetic distribution of all clades in the mammal-wide analysis across all viruses and within coronaviruses and flaviviruses for mean CFR, maximum CFR, and the percent of viruses with onward transmission. Of the five viral families, we only identified risky bat clades for the coronaviruses and flaviviruses. Clade numbers match those provided in Table S2.

In the virus-wide analysis, four clades differed in mean CFRs, and all were risky (Figure 2, Table S2). Phylogenetic factorization first identified a bat subclade consisting of the superfamilies Emballonuroidea and Vespertilionoidea (Table 1). The second was a subclade of 13 families of the order Carnivora (Table S2)^37^. The third was a subclade of many residual bat families from both the Yangochiroptera and Yinpterochiroptera: Rhinopomatidae, Megadermatidae, Rhinolophidae, Hipposideridae, Pteropodidae, Noctilionidae, Mormoopidae, and Phyllostomidae (Table 1). The fourth was a subclade of the rodent family Cricetidae (Table S2)^37^.

**Table 1.** Risky clades of bats identified by phylogenetic factorization for mean CFR (A), maximum CFR (B), and the percent of viruses with onward transmission (C) among all mammals, stratified across all viruses and within select virus families. Taxonomy matches our mammal phylogeny, and clades are presented with the number of included species and the mean value of the given response variable (clade) relative to that in the paraphyletic remainder (other).

|  | Virus | Factor | Taxa | Tips | Clade | Other |
| --- | --- | --- | --- | --- | --- | --- |
| (A) | All viruses | 1 | Natalidae, Molossidae, Vespertilionidae, Nycteridae, Emballonuridae | 109 | 0.69 | 0.19 |
|  | All viruses | 3 | Rhinopomatidae, Megadermatidae, Rhinolophidae, Hipposideridae, Pteropodidae, Noctilionidae, Mormoopidae, Phyllostomidae | 111 | 0.35 | 0.12 |
|  | *Coronaviridae* | 1 | *Scotophilus, Pipistrellus, Nyctalus, Vespertilio, Neoromicia, Ia* | 12 | 0.31 | 0.08 |
|  | *Flaviviridae* | 2 | Megadermatidae, Rhinolophidae, Hipposideridae | 12 | 0.22 | 0.06 |
|  | *Flaviviridae* | 4 | Vespertilionidae | 21 | 0.15 | 0.07 |
| (B) | *Coronaviridae* | 1 | *Scotophilus, Pipistrellus, Nyctalus, Vespertilio, Neoromicia, Ia* | 12 | 0.32 | 0.08 |
|  | *Flaviviridae* | 2 | Megadermatidae, Rhinolophidae, Hipposideridae, Pteropodidae, Noctilionidae, Mormoopidae, Phyllostomidae, Natalidae, Molossidae, Vespertilionidae, Emballonuridae | 75 | 0.16 | 0.07 |
| (C) | All viruses | 2 | Rhinopomatidae, Megadermatidae, Rhinolophidae, Hipposideridae, Pteropodidae | 63 | 0.49 | 0.26 |
|  | *Flaviviridae* | 2 | Megadermatidae, Rhinolophidae, Hipposideridae, Pteropodidae, Mormoopidae, Phyllostomidae, Natalidae, Molossidae, Vespertilionidae, Emballonuridae | 72 | 0.33 | 0.09 |

For maximum CFR across all viruses, phylogenetic factorization identified a risky subclade within the infraclass Eutheria (i.e., placental mammals)^37^. The second was “non-risky” (i.e., harboring unusually low maximum CFRs) and included families from the superorder Cetartiodactyla (Table S2)^37^. The third was risky and comprised genera from the rodent subfamily Neotominae (Table S2)^37^. The fourth was non-risky and included bat families from the superfamily Rhinolophoidae (i.e., Rhinopomatidae, Megadermatidae, Rhinolophidae, and Hipposideridae; Table S2)^17^. Finally, the fifth was also non-risky and included families from the Carnivora suborder Caniformia (Table S2)^37^.

For the fraction of viruses with onward transmission, all clades identified by phylogenetic factorization were risky. The first clade included the primate parvorder Catarrhini (i.e., apes and “Old World” monkeys; Table S2)^37^. The second was a subclade of the bat suborder Yinpterochiroptera, consisting of the superfamily Pteropodidae and the families Rhinopomatidae, Megadermatidae, Rhinolophidae, and Hipposideridae (Table 1)^17^. The third was a subclade from the rodent subfamily Sigmodonitae (Table S2)^37^. The fourth was composed of primate families from the infraorder Simiiformes and the Tupaiidae family from the order Scandentia (Table S2)^37^. Importantly, the order Chiroptera as a whole was not organically identified as a risky clade in any of these analyses.

When stratifying our mammal-wide analyses by the five selected virus families with sufficient sample sizes, phylogenetic factorization identified nine clades that differed in mean CFR, ten in maximum CFR, and four in the fraction of viruses with onward transmission. Within coronaviruses, the same risky bat clade was identified by phylogenetic factorization for mean CFR and maximum CFR: a subclade of the cosmopolitan subfamily Vespertilioninae (including only the genera *Scotophilus, Pipistrellus, Nyctalus, Vespertilio, Neoromicia*, and *Ia*, which are all only found in the eastern hemisphere)^17^. Within flaviviruses, two risky bat clades were identified for mean CFR: one included the bat families Megadermatidae, Rhinolophidae, and Hipposideridae (all restricted to the eastern hemisphere), and the other was the cosmopolitan family Vespertilionidae (Table 1, Figure 2)^17^. For maximum CFR, the one risky bat clade consisted of families in both the Yinpterochiroptera and Yangochiroptera (Table 1, Figure 2). For the fraction of viruses with onward transmission, one risky bat subclade was identified spanning the Yangochiroptera and Yinpterochiroptera, similar to clades identified for mean and maximum CFR (Table 1, Figure 2). For a full list of all clades identified, see Table S1. Overall, the bat clades identified by phylogenetic factorization were mostly similar between the mammal-wide and bat-specific analyses, both across viruses and within specific virus families (Tables S2 and S3). Importantly, analyses within each of our five viral families only found risky clades for coronaviruses and flaviviruses.

### Geographic hotspots of “risky” bat clades

We next extracted the members of risky bat clades identified by phylogenetic factorization in the mean CFR analysis and mapped their geographic distributions with anthropogenic footprint (Figure 3). Across all viruses, geographic hotspots of zoonotic risk occurred in Central America, coastal South America, equatorial Africa, and Southeast Asia (Figure 3). In the coronavirus analysis, hotspots clustered in Europe, as well as Sub-Saharan Africa and East Asia (Figure 3). In the flavivirus analysis, hotspots occurred in the Americas, Europe, Southeast Asia, and Sub-Saharan Africa (Figure 3).

**Figure 3.**
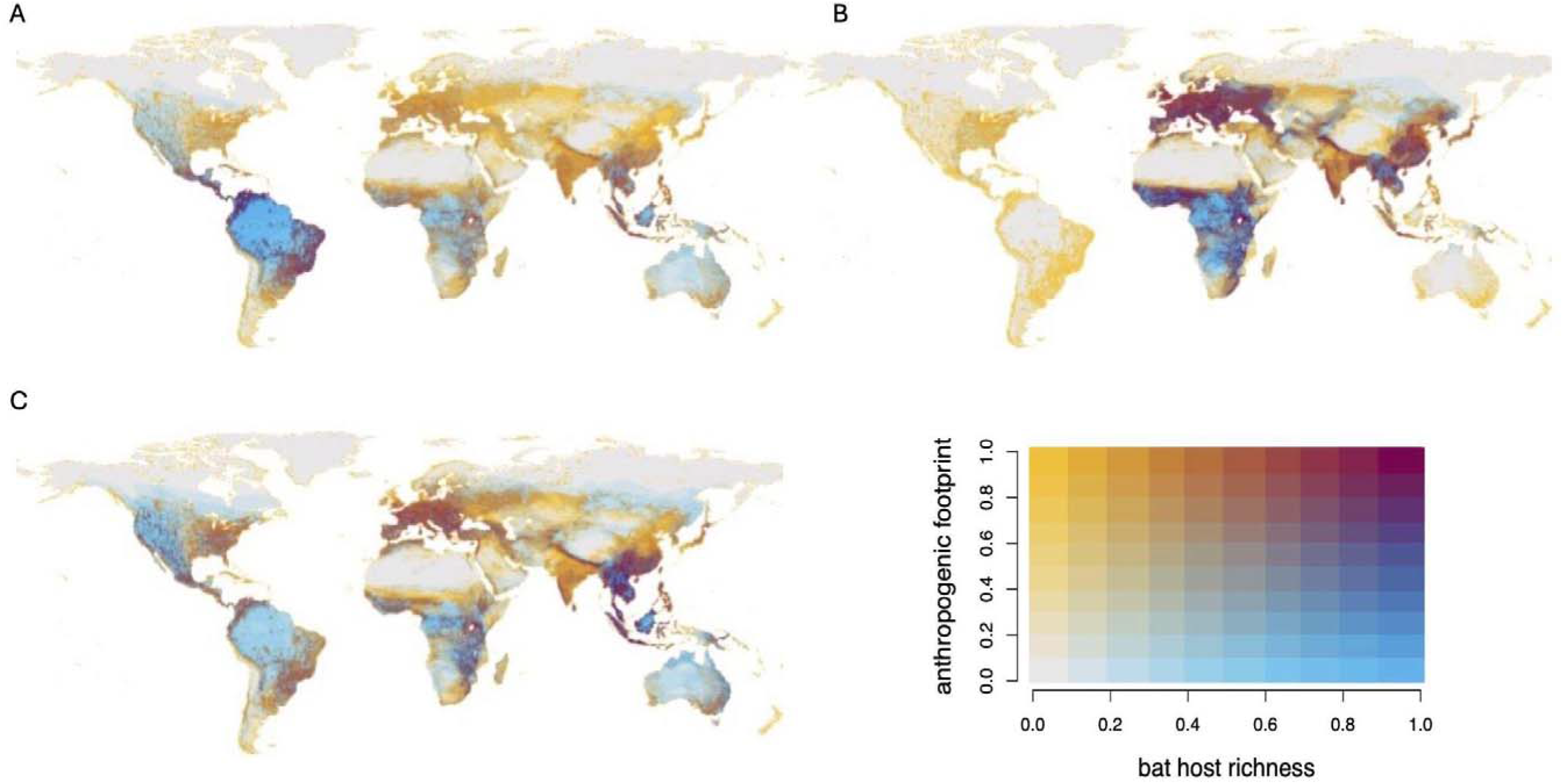
Geographic distribution of risky bat clades in the mammal-wide analysis (a) across all viruses and within (b) coronaviruses and (c) flaviviruses for mean CFR, co-mapped with anthropogenic footprint. Of the five viral families, we only identified risky bat clades for the coronaviruses and flaviviruses. Red highlights regions to prioritize for viral surveillance and ecological interventions due to increased spillover risk.

## Discussion

Zoonotic pathogens from wildlife have caused many pandemics and represent the greatest number of high-impact emerging infectious diseases ^38^. Prior research states bats host high viral diversity and the greatest number of viruses with high virulence in humans ^8,14,39^. We investigated whether bats, as an entire order, emerge organically to have a higher propensity to host virulent viruses than other taxa or if analyses that are agnostic to taxonomic order would instead identify subclades of bats exhibiting high virulence. Using several approaches to assess phylogenetic patterns, our analyses show that bats do not come out as a group with uniform viral epidemic potential: virulence and transmissibility are clustered only within distinct subclades of bats, often within cosmopolitan families spanning both the western and eastern hemispheres.

In the bat subclades, we identified species across all viruses and within two viral families, the *Coronaviridae* and *Flaviviridae*, that are more likely than other groups of this order to host virulent and transmissible viruses. These results add further nuance to the discussion regarding bats as viral hosts: in lieu of possessing special immune systems, each bat species may possess unique immunological adaptations to harbor the particular viruses with which they have coevolved (like all mammals), and many of these traits exist due to the hyperdiverse nature of the Chiropteran order ^11,40–43^.

However, our results also raise the question of whether all bats uniformly tolerate highly virulent viruses, in addition to differentially harboring viruses ^12,44–46^. Tolerance describes an individual coexisting with a replicating virus and exhibiting few to no clinical signs of infection ^11,47^. Parsing whether species both carry and tolerate viruses will provide further insight into whether bats possess exceptional immune systems and can inform spillover prevention strategies ^11,47^. Future experimental viral infection studies could compare the immune responses and infection outcomes between bat species from the risky clades identified here and taxonomically relevant outgroups to the same virus to begin investigating if risky species exhibit dissimilar abilities to tolerate viruses ^40,48^.

Our results align with previous work showing certain virus families exhibit non-random associations with specific kinds of host species ^4–6,49,50^. Phylogenetic signal was notably high for coronaviruses, and a small subclade of the Vespertilioninae subfamily factored out with especially high virulence. This suggests that closely related bat species within this clade possess coronaviruses with similar virulence in humans. Any bat in the genera *Scotophilus* (*n* = 20), *Pipistrellus* (*n* = 33), *Nyctalus* (*n* = 8), *Vespertilio* (*n* = 2), *Neoromicia* (*n* = 6), and *Ia* (*n* = 1) should, on average, be more likely to harbor coronaviruses with high virulence in humans ^17^. As surveillance across the Chiroptera is not cost-, labor-, or time-effective due to bats being extremely speciose and challenging to sample (e.g., nocturnal lifestyle, remote roosts), surveillance could monitor these species to discover novel hosts and coronaviruses ^4^. Further, while these bats are part of the cosmopolitan Vespertilionidae family, the subclade factored out only resides in the eastern hemisphere, so habitats where overlap of risky species and bat–human interactions are high could be prioritized both for surveillance and conservation to minimize negative human–bat interactions, providing the two-fold benefit of reducing spillover risk of high-risk viruses and conferring conservation benefits to bats, other wildlife, and humans ^9,51,52^.

A key finding from our pan-virus and *Flaviviridae* analyses is the bat superfamilies identified to harbor more virulent viruses than other mammals (and other bats) largely have cosmopolitan distributions. In the pan-virus analysis, this phenomenon may stem from bat–virus coevolution: viruses typically coevolve with their hosts either by specialization (e.g., recombination) or diversification (e.g., host switching)^42^. These data suggest diversification was selected for, facilitating future host jumps ^42^. As cosmopolitan families possess broad geographic distributions, diversification is likely advantageous for viruses as cosmopolitan species could introduce the viruses they carry to many potential hosts ^42^.

In the *Flaviviridae* analysis, virulent flaviviruses may cluster within cosmopolitan bats due to the nature of flaviviruses ^53^. All flaviviruses in this analysis were vector-borne, meaning they are transmitted between hosts via biting arthropods. Vector-borne viruses are more likely to be virulent because these viruses do not possess the evolutionary constraint of needing to keep their host alive for onward transmission ^53^. Alternatively, bats with cosmopolitan distributions may be more likely to carry virulent flaviviruses simply because they are exposed to a higher diversity of flaviviruses, owing to their broad distributions and thus greater exposure risk ^54,55^.

Of note, whether bats play an important role in the transmission of flaviviruses is unclear, as our understanding of viral replication of flaviviruses within bats is limited ^56,57^. When host-virus associations via a vector were removed from the data, zero species of bats were associated with flaviviruses and togaviruses, meaning these viruses may not replicate in bats, key for onward transmission ^57^. We, therefore, caution how the results of our analyses are interpreted as not all species may play a direct role in the circulation of these viruses ^57^. That being said, the role of bats and other hosts in the transmission of most viruses has yet to be fully understood^56,57^.

Our final analysis focused on geographic distributions of bat species found to harbor viruses with high CFRs. Those members of the risky clades identified here not included in our analysis could be prioritized for viral discovery, and the overlap with anthropogenic footprint can provide targeted areas for detection of viruses due to increased potential of spillover into humans ^51^. For example, *Nyctalus lasiopterus* (present across Eastern Europe, France, and Spain) and *Pipistrellus javanicus* (found throughout Southeast Asia) from the high-virulence clade in the coronavirus analysis are not known to host coronaviruses but would be good targets because their habitats exhibit high anthropogenic pressure ^4^. These two species have also been identified in previous machine learning analyses as high-priority targets for coronavirus surveillance due to their similar ecologies to known hosts.^4^ As human activities expand, better understanding where and how anthropogenic footprint affects viral dynamics is crucial for quantifying future spillover risk to humans and amplifying hosts, like domestic animals and threatened wildlife (e.g., primates)^51,58^. For example, deforestation is connected with increased Nipah virus spillovers from bats to pigs and humans, and the loss of winter habitat for *Pteropus* bats in Australia has increased Hendra virus spillovers from these animals ^51,59^.

Our phylogenetic analyses do not *a priori* assume a particular phylogenetic scale, which can help uncover particular clades of interest above or below a typically specified taxon (e.g., mammalian order) ^22,60^. They can also complement trait-based analyses and help guide new research directions to explain particular phylogenetic clustering ^22^. Our results can encourage next identifying the underlying traits that differentiate bat clades with high virulence viruses from other bat clades ^50^. For example, bats exhibit diverse immune systems and use different (and often non-overlapping) adaptations to cope with viruses (e.g., constitutive expression of interferons, increased heat shock protein expression, dampened inflammatory response) ^11,43^. Since selection for virulence is influenced by the host immune system, comparing potential dissimilarities between the immune systems of risky and non-risky clades of bats will be an important area for future work ^42,61,62^.

Finally, bats perform critical ecosystem services, like seed dispersal, pollination, and nutrient redistribution ^18^. Bats in the cosmopolitan families Molossidae, Vespertilionidae, and Emballonuridae are either obligate or facultative insectivores, consuming insects known to be agricultural pests ^18^. Bats also provide recreational and cultural value to humans and inspire technological innovations ^18,63,64^. However, many species are at risk of extinction due to habitat loss and climate change, and these risks are exacerbated by bat exterminations driven by fears over perceptions of bats-as-virus-hosts ^18,19^. For the latter point, culling bat colonies can have unintended consequences, as it can increase viral prevalence in bats and, in turn, amplify spillover risk ^51,65^. We hope our analyses can aid in facilitating conversations highlighting the ways in which human activities, not bats inherently, drive zoonotic viral emergence.^1,51^

## Acknowledgements

We thank members of the Becker Lab at the University of Oklahoma as well as Cole Brookson and Ricardo Rivero for helpful feedback.

## Data Availability

Viral virulence data and the mammalian phylogeny are available from previous publications ^5,8,25^ Mammal–virus associations are available through VIRION ^23^ All R code to reproduce analyses is available in the project GitHub repository: https://github.com/viralemergence/phylofatality.

