## Supplementary material for "Viral epidemic potential is not uniformly distributed across the bat phylogeny": Supplmental Information

Table S1. Tests of phylogenetic signal in mean CFR (A), maximum CFR (B), and fraction of viruses with onward transmission (C) across all mammals and bats specifically, stratified across all viruses and within select virus families. Estimates of λ are provided with 95% confidence intervals alongside *p* values from tests of phylogenetic randomness (*p*_random_) and Brownian motion models of evolution (*p_BM_*).

|  | **All mammals** | | | | | **Bats** | | | | |
| --- | --- | --- | --- | --- | --- | --- | --- | --- | --- | --- |
|  | **Virus** | **λ** | **CI** | ***p_random_*** | ***p_BM_*** | **Virus** | **λ** | **CI** | ***p_random_*** | ***p_BM_*** |
| (A) | All viruses | 0.69 | 0.58–0.78 | <0.001 | <0.001 | All viruses | 0.87 | 0.69–0.95 | <0.001 | <0.001 |
|  | *Coronaviridae* | 1.00 | 0.99–1.00 | <0.001 | 1 | Coronaviruses | 1.00 | 0.99–1.00 | <0.001 | 1 |
|  | *Flaviviridae* | 0.79 | 0.65–0.89 | <0.001 | <0.001 | Flaviviruses | 0.97 | 0.65–1.00 | <0.001 | <0.001 |
|  | *Rhabdoviridae* | 0.72 | 0.57–0.83 | <0.001 | <0.001 | Rhabdoviruses | 0.52 | 0.23–0.80 | <0.001 | <0.001 |
|  | *Togaviridae* | 0.71 | 0.39–0.89 | <0.001 | <0.001 | Togaviruses | 0.50 | 0–0.95 | 0.81 | <0.001 |
|  | *Paramyxoviridae* | 0.00 | 0–0.72 | 1 | <0.001 | Paramyxoviruses | 0.00 | 0–0.51 | 1 | <0.001 |
| (B) | All viruses | 0.64 | 0.53–0.73 | <0.001 | <0.001 | All viruses | 0.71 | 0.40–0.87 | <0.001 | <0.001 |
|  | *Coronaviridae* | 1.00 | 0.99–1.00 | <0.001 | 1 | Coronaviruses | 1.00 | 0.99–1.00 | <0.001 | 1 |
|  | *Flaviviridae* | 0.67 | 0.51–0.79 | <0.001 | <0.001 | Flaviviruses | 0.00 | 0–0.33 | 1 | <0.001 |
|  | *Rhabdoviridae* | 0.87 | 0.77–0.93 | <0.001 | <0.001 | Rhabdoviruses | 0.00 | 0–1.00 | 1 | 1 |
|  | *Togaviridae* | 0.61 | 0.36–0.80 | <0.001 | <0.001 | Togaviruses | 0.00 | 0–0.63 | 1 | <0.001 |
|  | *Paramyxoviridae* | 0.89 | 0.33–0.98 | 0.03 | <0.001 | Paramyxoviruses | 0.00 | 0–0.94 | 1 | <0.001 |
| (C) | All viruses | 0.85 | 0.80–0.90 | <0.001 | <0.001 | All viruses | 0.81 | 0.60–0.91 | <0.001 | <0.001 |
|  | *Flaviviridae* | 0.70 | 0.56–0.81 | <0.001 | <0.001 | Flaviviruses | 0.52 | 0.09–0.82 | 0.03 | <0.001 |
|  | *Togaviridae* | 1.00 | 1.00–1.00 | <0.001 | 1 | Togaviruses | 1.00 | 0.99–1.00 | <0.001 | 1 |
|  | *Paramyxoviridae* | 0.00 | 0–0.86 | 1 | <0.001 | Paramyxoviruses | 0.96 | 0–1.00 | 0.26 | <0.001 |

Table S2. Phylogenetic factorization for mean CFR (A), maximum CFR (B), and fraction of viruses with onward transmission (C) across all mammals, stratified across all viruses and within select virus families. Taxonomy matches our mammal phylogeny, and clades are presented with the number of included species and the mean value of the given response variable (clade) relative to that in the paraphyletic remainder (other).

|  | **Virus** | **Factor** | **Taxa** | **Tips** | **Clade** | **Other** |
| --- | --- | --- | --- | --- | --- | --- |
| (A) | All viruses | 1 | Natalidae, Molossidae, Vespertilionidae, Nycteridae, Emballonuridae | 109 | 0.69 | 0.19 |
|  | All viruses | 2 | Canidae, Phocidae, Odobenidae, Otariidae, Mephitidae, Procyonidae, Mustelidae, Ailuridae, Ursidae, Hyaenidae, Herpestidae, Viverridae, Felidae | 114 | 0.43 | 0.15 |
|  | All viruses | 3 | Rhinopomatidae, Megadermatidae, Rhinolophidae, Hipposideridae, Pteropodidae, Noctilionidae, Mormoopidae, Phyllostomidae | 111 | 0.35 | 0.12 |
|  | All viruses | 4 | *Neotoma, Reithrodontomys, Onychomys, Peromyscus, Megadontomys, Baiomys, Tylomys, Sigmodon, Bibimys, Necromys, Akodon, Thaptomys, Oxymycterus, Zygodontomys, Hylaeamys, Nephelomys, Oecomys, Transandinomys, Handleyomys, Neacomys, Oligoryzomys, Nectomys, Melanomys, Oryzomys, Sooretamys, Calomys, Phyllotis, Loxodontomys, Graomys, Abrothrix* | 83 | 0.31 | 0.09 |
|  | *Coronaviridae* | 1 | *Scotophilus, Pipistrellus, Nyctalus, Vespertilio, Neoromicia, Ia* | 12 | 0.31 | 0.08 |
|  | *Coronaviridae* | 2 | Equidae, Rhinocerotidae, Camelidae, Suidae, Iniidae, Phocoenidae, Monodontidae, Delphinidae, Bovidae | 16 | 0.15 | 0.06 |
|  | *Flaviviridae* | 1 | Pitheciidae, Callitrichidae, Cebidae, Aotidae, Atelidae, Hylobatidae, Hominidae, Cercopithecidae | 60 | 0.22 | 0.07 |
|  | *Flaviviridae* | 2 | Megadermatidae, Rhinolophidae, Hipposideridae | 12 | 0.22 | 0.06 |
|  | *Flaviviridae* | 3 | Tragulidae, Giraffidae, Antilocapridae, Cervidae, Bovidae | 60 | 0.01 | 0.07 |
|  | *Flaviviridae* | 4 | Vespertilionidae | 21 | 0.15 | 0.07 |
|  | *Rhabdoviridae* | 1 | Canidae, Mephitidae, Procyonidae, Mustelidae, Ursidae, Hyaenidae, Herpestidae, Viverridae, Felidae, Equidae, Camelidae, Tayassuidae, Suidae, Antilocapridae, Cervidae, Bovidae, Rhinolophidae, Hipposideridae, Pteropodidae, Mormoopidae, Phyllostomidae, Molossidae, Vespertilionidae, Nycteridae, Emballonuridae | 238 | 0.9 | 0.38 |
|  | *Rhabdoviridae* | 2 | Sciuridae | 10 | 0.93 | 0.29 |
|  | *Togaviridae* | 1 | Canidae, Phocidae, Mephitidae, Procyonidae, Ursidae, Felidae, Equidae, Tapiridae, Camelidae, Tayassuidae, Suidae, Cervidae, Bovidae, Hipposideridae, Pteropodidae, Phyllostomidae, Molossidae, Vespertilionidae, Emballonuridae | 60 | 0.12 | 0.04 |
| (B) | All viruses | 1 | Manidae, Canidae, Phocidae, Odobenidae, Otariidae, Mephitidae, Procyonidae, Mustelidae, Ailuridae, Ursidae, Hyaenidae, Herpestidae, Viverridae, Felidae, Equidae, Tapiridae, Rhinocerotidae, Camelidae, Tayassuidae, Suidae, Hippopotamidae, Physeteridae, Iniidae, Phocoenidae, Monodontidae, Delphinidae, Eschrichtiidae, Balaenopteridae, Balaenidae, Tragulidae, Giraffidae, Antilocapridae, Cervidae, Bovidae, Rhinopomatidae, Megadermatidae, Rhinolophidae, Hipposideridae, Pteropodidae, Noctilionidae, Mormoopidae, Phyllostomidae, Natalidae, Molossidae, Vespertilionidae, Nycteridae, Emballonuridae | 468 | 0.57 | 0.24 |
|  | All viruses | 2 | Tayassuidae, Suidae, Hippopotamidae, Physeteridae, Iniidae, Phocoenidae, Monodontidae, Delphinidae, Eschrichtiidae, Balaenopteridae, Balaenidae, Tragulidae, Giraffidae, Antilocapridae, Cervidae, Bovidae | 118 | 0.22 | 0.69 |
|  | All viruses | 3 | *Neotoma, Reithrodontomys, Onychomys, Peromyscus, Megadontomys, Baiomys* | 30 | 0.72 | 0.2 |
|  | All viruses | 4 | Rhinopomatidae, Megadermatidae, Rhinolophidae, Hipposideridae | 30 | 0.33 | 0.72 |
|  | All viruses | 5 | Phocidae, Odobenidae, Otariidae | 16 | 0.21 | 0.75 |
|  | *Coronaviridae* | 1 | *Scotophilus, Pipistrellus, Nyctalus, Vespertilio, Neoromicia, Ia* | 12 | 0.32 | 0.08 |
|  | *Coronaviridae* | 2 | Equidae, Rhinocerotidae, Camelidae, Suidae, Iniidae, Phocoenidae, Monodontidae, Delphinidae, Bovidae | 16 | 0.15 | 0.06 |
|  | *Flaviviridae* | 1 | Pitheciidae, Callitrichidae, Cebidae, Aotidae, Atelidae, Hylobatidae, Hominidae, Cercopithecidae | 60 | 0.28 | 0.09 |
|  | *Flaviviridae* | 2 | Megadermatidae, Rhinolophidae, Hipposideridae, Pteropodidae, Noctilionidae, Mormoopidae, Phyllostomidae, Natalidae, Molossidae, Vespertilionidae, Emballonuridae | 75 | 0.16 | 0.07 |
|  | *Flaviviridae* | 3 | Elephantidae, Dasypodidae, Myrmecophagidae, Cyclopedidae, Megalonychidae, Bradypodidae | 10 | 0.18 | 0.07 |
|  | *Flaviviridae* | 4 | Canidae, Phocidae, Mephitidae, Procyonidae, Mustelidae, Ailuridae, Ursidae, Herpestidae, Felidae | 26 | 0.12 | 0.06 |
|  | *Flaviviridae* | 5 | Tragulidae, Giraffidae, Antilocapridae, Cervidae, Bovidae | 60 | 0.03 | 0.07 |
|  | *Rhabdoviridae* | 1 | Canidae, Mephitidae, Procyonidae, Mustelidae, Ursidae, Hyaenidae, Herpestidae, Viverridae, Felidae, Equidae, Camelidae, Tayassuidae, Suidae, Antilocapridae, Cervidae, Bovidae, Rhinolophidae, Hipposideridae, Pteropodidae, Mormoopidae, Phyllostomidae, Molossidae, Vespertilionidae, Nycteridae, Emballonuridae | 238 | 0.96 | 0.43 |
|  | *Rhabdoviridae* | 2 | Sciuridae | 10 | 1 | 0.32 |
|  | *Togaviridae* | 1 | Canidae, Phocidae, Mephitidae, Procyonidae, Ursidae, Felidae, Equidae, Tapiridae, Camelidae, Tayassuidae, Suidae, Cervidae, Bovidae, Hipposideridae, Pteropodidae, Phyllostomidae, Molossidae, Vespertilionidae, Emballonuridae | 60 | 0.17 | 0.06 |
| (C) | All viruses | 1 | Hylobatidae, Hominidae, Cercopithecidae | 81 | 0.83 | 0.28 |
|  | All viruses | 2 | Rhinopomatidae, Megadermatidae, Rhinolophidae, Hipposideridae, Pteropodidae | 63 | 0.49 | 0.26 |
|  | All viruses | 3 | *Bibimys, Necromys, Akodon, Thaptomys, Oxymycterus* | 16 | 0.86 | 0.25 |
|  | All viruses | 4 | Tupaiidae, Galagidae, Lorisidae, Indriidae, Lemuridae, Lepilemuridae, Pitheciidae, Callitrichidae, Cebidae, Aotidae, Atelidae | 52 | 0.51 | 0.24 |
|  | *Flaviviridae* | 1 | Pitheciidae, Callitrichidae, Cebidae, Aotidae, Atelidae, Hylobatidae, Hominidae, Cercopithecidae | 60 | 0.64 | 0.16 |
|  | *Flaviviridae* | 2 | Megadermatidae, Rhinolophidae, Hipposideridae, Pteropodidae, Mormoopidae, Phyllostomidae, Natalidae, Molossidae, Vespertilionidae, Emballonuridae | 72 | 0.33 | 0.09 |
|  | *Flaviviridae* | 3 | Talpidae, Soricidae, Erinaceidae, Canidae, Phocidae, Mephitidae, Procyonidae, Mustelidae, Ailuridae, Ursidae, Herpestidae, Felidae, Equidae, Rhinocerotidae, Camelidae, Tayassuidae, Suidae, Delphinidae, Giraffidae, Antilocapridae, Cervidae, Bovidae, Galagidae, Lorisidae, Indriidae, Lemuridae, Lepilemuridae, Ochotonidae, Leporidae, Gliridae, Sciuridae, Ctenodactylidae, Hystricidae, Dasyproctidae, Caviidae, Erethizontidae, Echimyidae, Dipodidae, Cricetidae, Muridae, Heteromyidae, Geomyidae | 180 | 0.06 | 0.36 |
|  | *Togaviridae* | 1 | Galagidae, Lemuridae, Cebidae, Atelidae, Hominidae, Cercopithecidae | 17 | 0.83 | 0.05 |

Table S3. Phylogenetic factorization for mean CFR (A), maximum CFR (B), and fraction of viruses with onward transmission (C) within only bats, stratified across all viruses and within select virus families. Taxonomy matches our mammal phylogeny, and clades are presented with the number of included species and the mean value of the given response variable (clade) relative to that in the paraphyletic remainder (other).

|  | **Virus** | **Factor** | **Taxa** | **Tips** | **Clade** | **Other** |
| --- | --- | --- | --- | --- | --- | --- |
| (A) | All viruses | 1 | Natalidae, Molossidae, Vespertilionidae, Nycteridae, Emballonuridae | 109 | 0.67 | 0.37 |
|  | All viruses | 2 | Rhinopomatidae, Megadermatidae, Rhinolophidae, Hipposideridae | 30 | 0.14 | 0.43 |
|  | *Coronaviridae* | 1 | *Scotophilus, Pipistrellus, Nyctalus, Vespertilio, Neoromicia, Ia* | 12 | 0.31 | 0.06 |
|  | *Flaviviridae* | 1 | Megadermatidae, Rhinolophidae, Hipposideridae | 12 | 0.21 | 0.09 |
|  | *Flaviviridae* | 2 | Vespertilionidae | 21 | 0.15 | 0.06 |
|  | *Rhabdoviridae* | 1 | *Lonchophylla, Artibeus, Dermanura, Vampyrodes, Platyrrhinus, Uroderma, Sturnira, Carollia* | 12 | 0.65 | 0.97 |
|  | *Rhabdoviridae* | 2 | *Diaemus, Desmodus, Diphylla, Micronycteris, Lonchorhina, Glossophaga, Leptonycteris, Anoura, Trachops, Phyllostomus, Lophostoma, Chrotopterus* | 17 | 0.87 | 1 |
| (B) | All viruses | 1 | Rhinopomatidae, Megadermatidae, Rhinolophidae, Hipposideridae | 30 | 0.32 | 0.76 |
|  | All viruses | 2 | *Lonchophylla, Artibeus, Dermanura, Vampyrodes, Platyrrhinus, Uroderma, Sturnira, Rhinophylla, Carollia, Trachops, Phyllostomus, Tonatia, Lophostoma, Chrotopterus* | 29 | 0.48 | 0.81 |
|  | *Coronaviridae* | 1 | *Scotophilus, Pipistrellus, Nyctalus, Vespertilio, Neoromicia, Ia* | 12 | 0.32 | 0.06 |
|  | *Flaviviridae* | 1 | Megadermatidae, Rhinolophidae, Hipposideridae | 12 | 0.27 | 0.15 |
|  | *Rhabdoviridae* | 1 | *Lonchophylla, Artibeus, Dermanura, Vampyrodes, Platyrrhinus, Uroderma, Sturnira, Carollia, Trachops, Phyllostomus, Lophostoma, Chrotopterus* | 17 | 0.83 | 1 |
| (C) | All viruses | 1 | Mormoopidae, Phyllostomidae, Natalidae, Molossidae, Vespertilionidae, Nycteridae, Emballonuridae | 153 | 0.24 | 0.49 |
